# Modeling the role of the thalamus in resting-state functional connectivity: nature or structure

**DOI:** 10.1101/2023.03.07.531650

**Authors:** Jesús Cabrera-Álvarez, Nina Doorn, Fernando Maestú, Gianluca Susi

## Abstract

The thalamus is a central brain structure that serves as a relay station for sensory inputs from the periphery to the cortex and regulates cortical arousal. Traditionally, it has been regarded as a passive relay that transmits information between brain regions. However, recent studies have suggested that the thalamus may also play a role in shaping functional connectivity (FC) in a task-based context. Based on this idea, we hypothesized that due to its centrality in the network and its involvement in cortical activation, the thalamus may also contribute to resting-state FC, a key neurological biomarker widely used to characterize brain function in health and disease. To investigate this hypothesis, we constructed ten in-silico brain network models based on neuroimaging data (MEG, MRI, and dwMRI), and simulated them including and excluding the thalamus. and raising the noise into thalamus to represent the afferences related to the reticular activating system (RAS) and the relay of peripheral sensory inputs. We simulated brain activity and compared the resulting FC to their empirical MEG counterparts to evaluate model’s performance. Results showed that a parceled version of the thalamus with higher noise, able to drive damped cortical oscillators, enhanced the match to empirical FC. However, with an already active self-oscillatory cortex, no impact on the dynamics was observed when introducing the thalamus. We also demonstrated that the enhanced performance was not related to the structural connectivity of the thalamus, but to its higher noisy inputs. Additionally, we highlighted the relevance of a balanced signal-to-noise ratio in thalamus to allow it to propagate its own dynamics. In conclusion, our study sheds light on the role of the thalamus in shaping brain dynamics and FC in resting-state and allowed us to discuss the general role of criticality in the brain at the mesoscale level.

**Author summary:** Synchrony between brain regions is an essential aspect of coordinated brain function and serves as a biomarker of health and disease. The thalamus, due to its centrality and widespread connectivity with the cortex, is a crucial structure that may contribute to this synchrony by allowing distant brain regions to work together. In this study, we used computational models to investigate the thalamus’s role in generating brain synchrony at rest. Our findings suggest that the structural connectivity of the thalamus is not its primary contribution to brain synchrony. Instead, we found that the thalamus plays a critical role in driving cortical activity, and when it is not driving this activity, its impact on brain synchrony is null. Our study provides valuable insights into the thalamocortical network’s role in shaping brain dynamics and FC in resting state, laying the groundwork for further research in this area.

## Introduction

In humans, the thalamus is a nut size structure near the center of the brain that relays sensory inputs traveling to the cortex [1], fosters cortico-cortical communication through transthalamic pathways [2, 3], and controls cortical arousal through the reticular activating system (RAS) [4]. To carry out these tasks, it contains three functionally distinct parts [5]: dorsal, ventral, and intralaminar. The dorsal part communicates bidirectionally with the cortex establishing two schemes of information exchange [6, 7] (see Figure 1): first-order relay, in which the thalamus receives subcortical and sensory inputs (i.e., driving inputs), relays them to the cortex through excitatory thalamocortical cells and gets back modulatory feedback from layer 6 pyramidal neurons (i.e., modulatory inputs); and higher order relay, in which the thalamus receives inputs from layer 5 pyramidal cells of a cortical region and relays them to another location in the cortex, creating a transthalamic pathway for cortico-cortical connections [2, 3]. In the ventral part, the reticular nucleus’ inhibitory neurons establish connections both with each other and with neurons in the dorsal nuclei, to regulate and foster communication inside the thalamus [8, 9]. The interactions between the dorsal and ventral parts of the thalamus allow for the generation of sustainable oscillations of neural activity, such as delta oscillations, sleep spindles, and slow waves, that may propagate to the cortex and influence its dynamics [10–15]. The intralaminar part is involved in the RAS [16] delivering cholinergic and monoaminergic neurotransmission diffusely to the cortex and controlling arousal [4].

**Fig 1.**
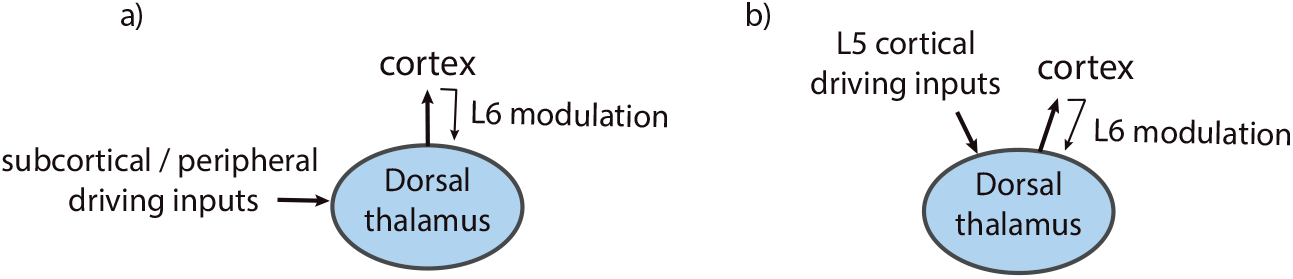
Main schemes of thalamocortical interaction. a) First-order relays, and b) higher-order relays

Through these pathways, the thalamus connects to a widespread set of cortical regions [17–19], playing a role in different psychological processes such as sleep [10, 20, 21], pain [22, 23], memory and learning [24], attention [25–27], motor and sensory processing [9, 15, 28], and consciousness [29–33]. This role of the thalamus has been classically depicted as a passive relay that transfers information between brain regions [24, 34], however recent findings are challenging this view [19, 24, 35–37]. Specifically, Schmitt et al. [25] showed that the mediodorsal thalamus was amplifying a FC pattern supporting the representation of specific task-related rules. All the thalamocortical mechanisms described above (i.e., the dorsal relay of peripheral sensory inputs, the transthalamic pathways for cortico-cortical communication, and the RAS neurotransmission that activates the cortex) may contribute to define the FC in the brain. Our study is aimed at understanding how.

FC is defined as a correlation between spatially distant neurophysiological signals [38] representing the functional integration of psychological processes in distributed brain networks [39]. Early neuroimaging studies were focused on revealing the activity patterns underlying cognitive processes during task execution, using resting-state as a control condition [40]. However, later findings showed that the brain in resting-state has a rich intrinsic activity [41, 42] related to automatic and unconscious cognitive processing [43, 44]. Since then, resting-state FC (rsFC) has been used to characterize brain function in health and disease [45–47]. It also characterizes aging, where a general decrease in FC has been shown [48, 49] and relates to changes in cognitive performance [50]. Interestingly, some authors have proposed that changes in the thalamocortical network may contribute substantially to the disruptions in FC and cognition during aging [51, 52]. Understanding the mechanisms that underlie and control FC is an important research question that is still undisclosed, especially in aging. Here, we hypothesize that a similar thalamic mechanism that has been shown to be involved in defining FC during task execution [25] might also be active in resting-state.

Computational modeling allows for the generation of in-silico versions of real brains and personalized brain dynamics [53] employing brain network models (BNM). A BNM is based on: a structural connectivity (SC) network derived from diffusion-weighted MRI that captures how brain regions are wired together, and a set of neural mass models (NMM) that reproduce the electrophysiological dynamics of each brain region. A widely studied NMM is the Jansen-Rit (JR; [54]), a biologically-inspired model of a cortical column that implements excitatory and inhibitory subpopulations to produce oscillatory activity. This model shows a bifurcation over a parameter representing the strength of its inputs [55, 56] that will be used in our work to reproduce different modes of the thalamocortical interaction. In a system, a bifurcation occurs when a change in the value of a parameter (i.e., bifurcation parameter) produces a qualitative change in the behavior of the system. For the JR model, the bifurcation separates two different states: a fixed point state, where the model behaves as a damped oscillator (prebifurcation), and a limit cycle state, where the model autonomously oscillates (postbifurcation). These two states turned out to be relevant to understand our findings.

To investigate the potential contribution of the thalamus to rsFC, we built ten in-silico BNMs based on healthy subjects’ neuroimaging data (MEG, MRI, dwMRI) using JR NMMs. We simulated them using: 1) three SC versions (i.e., parceled thalamus, pTh; thalamus as a single node, Th; without thalamus, woTh) to explore the effect of both the parcellation and the mere presence of cortico-cortical transthalamic pathways and 2) implementing a higher noisy input to the thalamus to reproduce its participation in RAS and the presence of peripheral sensory relays. We compared the simulated FC and dynamical FC (dFC) to their empirical MEG counterparts to evaluate performance. Additionally, we performed further simulations to explore under which conditions the thalamus contributes to the rsFC, including a control experiment using the cortico-cerebellar network instead of the thalamocortical one, and a set of parameter explorations over the intrinsic thalamic oscillatory behavior and the magnitude of the implemented noise. Our results showed that a limited set of driving nodes leading cortical activity was a plausible scenario in rsFC, where the thalamus would play a major role due to its nature: involved in the RAS system, and projecting sensory relays. These results contribute to the understanding of the basic principles of whole-brain function in health and disease, and to enrich the current picture of criticality behavior in the brain.

## Results

### The thalamus impacts rsFC through its afferences

To explore the role of the thalamus in rsFC, we used two features of our in-silico BNMs: its structure, by simulating three different SC versions per subject (pTh, Th, woTh), and the NMMs noisy input by implementing higher than cortex thalamic noise (*η_th_*=[0.022, 2.2×10^-8^], *η_cx_*=[2.2×10^-8^]) to represent thalamic RAS system and peripheral first-order relays. We used the coupling parameter (*g*) to scale connectivity weights looking for the best match to empirical rsFC (i.e., the working point), as usual in whole-brain modeling [57–60]. Given that *g* acts as a bifurcation parameter, the simulated activity could be categorized into two regimes: prebifurcation, in which nodes operate as damped oscillators, and postbifurcation, in which nodes operate as autonomous oscillators. We simulated 60 seconds of brain activity per model and measured FC and dFC in the alpha band to compare to their empirical counterparts using Pearson’s correlation (r_PLV_) and Kolmogorov-Smirnov distance (KSD), respectively. We will show statistical comparisons using the best values of those metrics per subject and bifurcation side.

Results on r_PLV_ showed a significant impact for both the structure [F(2, 18)=191.77, 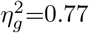, eps=0.55, p<0.0001], noise [F(1, 9)=178.25, 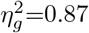, eps=1, p<0.0001], and their interaction [F(2, 18)=172.12, 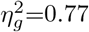, eps=0.55, p<0.0001] in *prebifurcation.* In contrast, in *postbifurcation,* we did not find significant differences for structure [F(2, 18)=1.29, 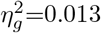, eps=0.76, p=0.29] or the interaction [F(2, 18)=1.74, 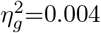, eps=0.65, p=0.21] while a weak effect was found for noise [F(1, 9)=5.89, 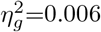, eps=1, p=0.038].

In prebifurcation, implementing high thalamic noise raised significantly r_PLV_ values from close to zero to r_PLV_≈0.33 for Th [W=0, Cohen’s d=5.09, p-corr=0.005], and r_PLV_≈0.45 for pTh [W=0, Cohen’s d=6.8, p-corr=0.005] (see Fig. 2). The resulting correlation values with high noise differed significantly between the implementations of the thalamic structure [F(2, 18)=186, 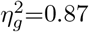, eps=0.53, p-corr<0.0001], and pTh showed a global peak of r_PLV_ that unexpectedly overcame the values observed in postbifurcation in 8 out of 10 subjects (see supplementary fig. S1 Fig) although the differences were not statistically significant [W=18, Cohen’s d=0.142, p=0.375].

**Fig 2.**
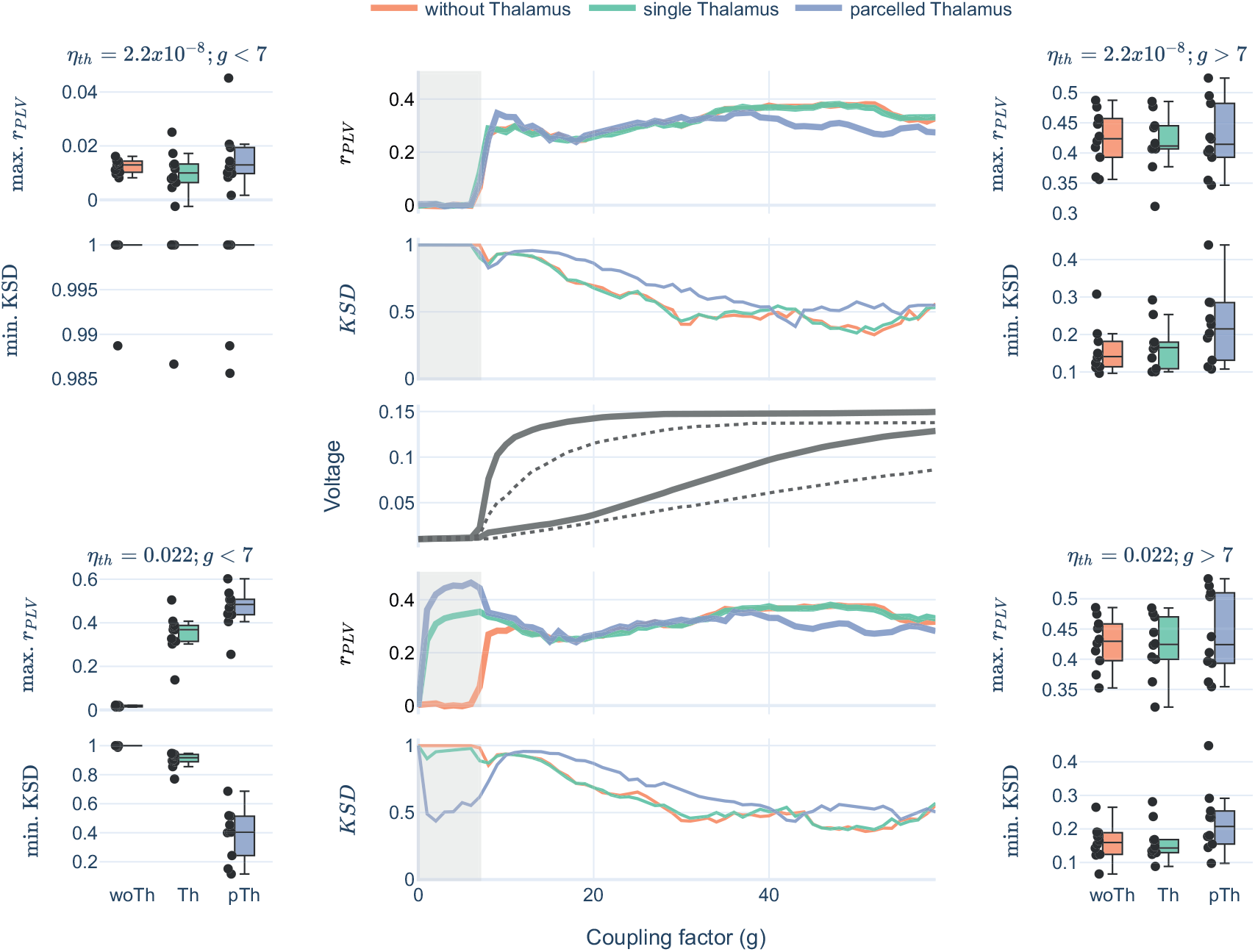
Thalamocortical experiment. Parameter space explorations over g, with two levels of thalamic noise and three SC versions. The central column shows averaged FC and dFC metrics with prebifurcation range shadowed. In the margins, boxplots for maximum values of r_PLV_ and minimum KSD per subject and thalamic structure. Central row showing bifurcation diagrams for the cortex (thick line) and the thalamus (dashed line) with maximum and minimum voltage per coupling factor.

Regarding dFC, the results followed a similar trend in which thalamic structure [F(2, 18)=119.95, 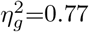, eps=0.56, p<0.0001], the thalamic noise [F(1, 9)=106.3, 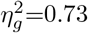, eps=1, p<0.0001] and the interaction [F(2, 18)=111.66, 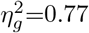, eps=0.55, p<0.0001] were statistically significant factors only in prebifurcation. In that range, high thalamic noise enhanced performance for pTh [W=0, Cohen’s d=4.89, p-corr=0.0029] showing a local KSD minimum, and also slightly for Th [W=0, Cohen’s d=2.5, p-corr=0.0029] (see Fig. 2). Interestingly, we noticed that high values of KSD for woTh and Th were due to opposite underlying dFC distributions. woTh correlations were centered near r=0 implying that FC matrices in time changed randomly, while Th correlations were centered near r=1 implying that FC matrices in time were quasistatic (see supplementary Fig. S2 Fig).

In summary, the inclusion of the thalamus in the model had an impact just in prebifurcation range (in which nodes operate as damped oscillators), and only when implementing a high thalamic noise. For pTh, this condition overcame the performance of any other model. In postbifurcation range (in which nodes autonomously oscillate) the values for r_PLV_ were also high, however, the thalamus did not show an impact.

### Structure is not the key: comparing thalamocortical and cortico-cerebellar networks

We wondered whether the observed improvement of r_PLV_ with high thalamic noise in prebifurcation could be explained by the specific characteristics of the SC pattern of the thalamus. To test this, we performed a control experiment comparing the thalamocortical network to another with similar properties: the cortico-cerebellar network (see Table 1). We built three SC versions per subject: parceled cerebellum (pCer), single node cerebellum (Cer) and without cerebellum (woCer). We simulated them implementing high noise into the cerebellum to compare the model performance to the thalamocortical network.

**Table 1.** Network features for the thalamus and the cerebellum.

|  | Degree | Node strength | Betweenness | Path Length |
| --- | --- | --- | --- | --- |
| Global average | 0.827 | 0.231 | 0.00119 | 1.165 |
| Thalamus average | 0.851 | 0.111 | 0.00125 | 1.141 |
| Cerebellum average | 0.883 | 0.224 | 0.00132 | 1.110 |

Averaged network metrics for the thalamus, the cerebellum, and the global average of all regions. Similar relation to average was observed between regions in all metrics (degree, betweenness, and path length) except for node strength. For more details on the network analysis, see S1 Table.

We observed that the general trend found with the thalamocortical networks persisted. woCer simulations showed close to zero r_PLV_ values in prebifurcation range with high noise, while Cer and pCer increased significantly their maximum correlations [F(2, 18)=148.39, 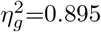, eps=0.75, p-corr<0.0001] up to similar values obtained with the thalamocortical network (see Fig 3). Moreover, pCer in prebifurcation also resulted in a global maximum in r_PLV_compared to postbifurcation values. Interestingly, Cer changed the underlying bifurcation diagram of the model, moving it toward higher values of coupling. This was reflected by the peak of r_PLV_ in higher g values.

**Fig 3.**
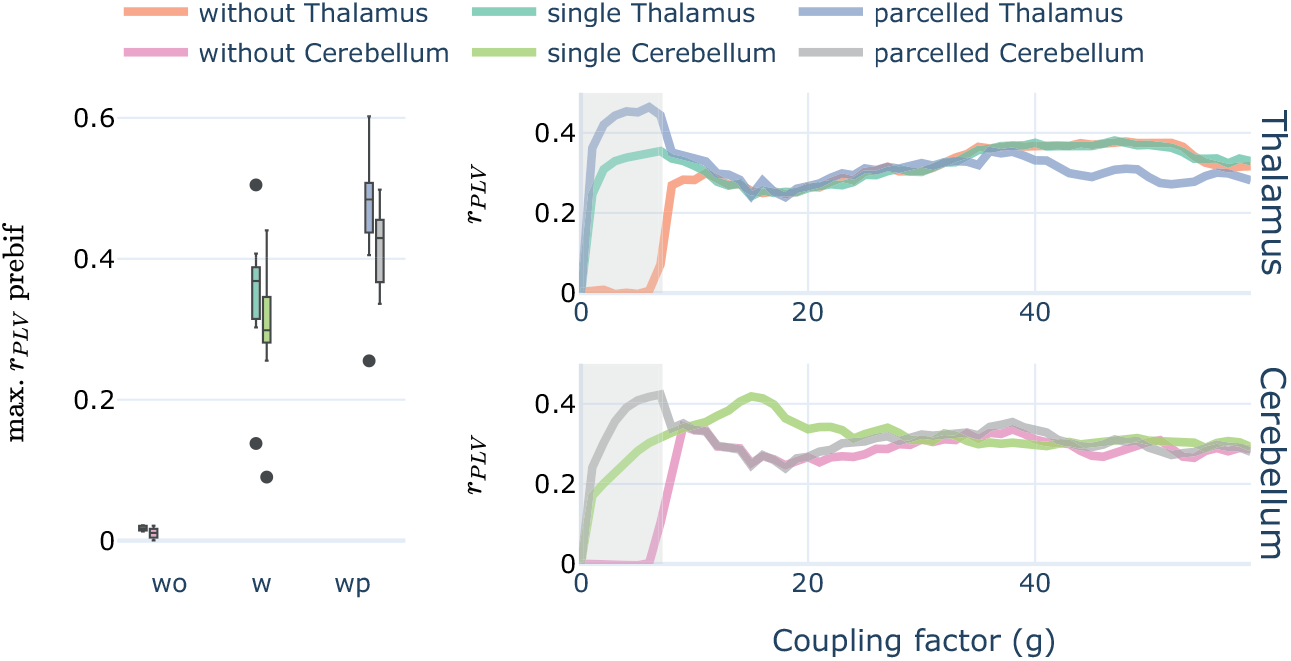
Cortico-cerebellar control experiment. Boxplot showing maximum r_PLV_ values per subject in the prebifurcation space, both for the experiments with the thalamus and the cerebellum. Right, lineplots conveying averaged parameter spaces per condition. The shadowed area covering the prebifurcation range.

These results suggest that the specific thalamocortical SC pattern is not a major determinant of the thalamic contribution to rsFC.

### Brain dynamics underlying each scenario

To understand the dynamics that underlie the observed values of r_PLV_ and KSD, we extracted a simulation sample per model condition with pTh (i.e., high/low thalamic noise, and pre- / post-bifurcation; see Fig 4).

**Fig 4.**
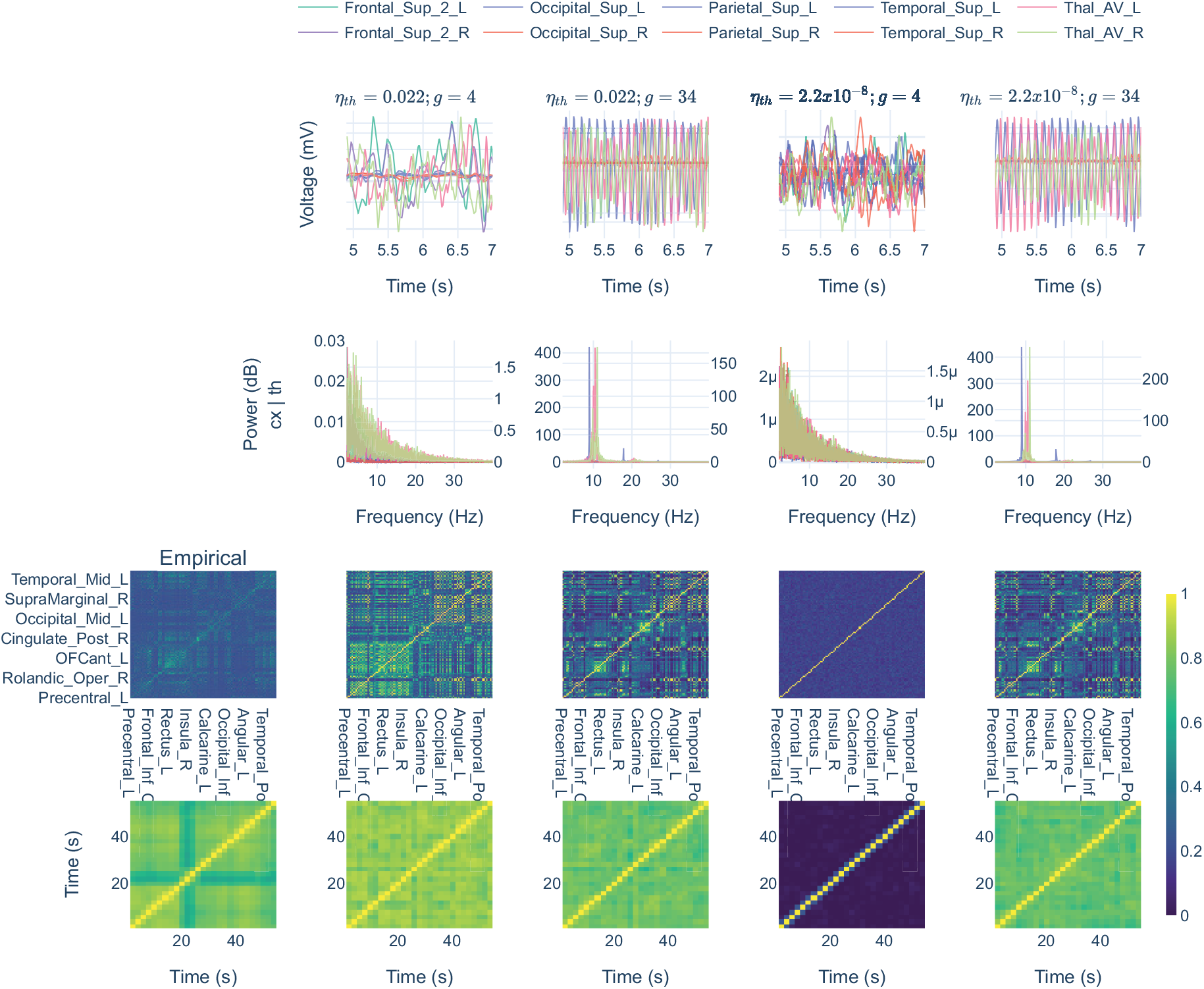
Simulation samples for one subject with parceled thalamus. Columns show one sample per simulation condition. Prebifurcation conditions were simulated with g=4, and postbifurcation was simulated with g=36. Rows showing demeaned signals, absolute spectra, PLV, and dFC matrices, respectively.

The dynamics in the prebifurcation range with high and low noise showed 1/f pink noise pattern. This is the result of a damped JR node processing a Gaussian noise as shown in previous literature [61, 62]. High and low noise conditions in prebifurcation could be differentiated by their spectral powers and by the differences in FC matrices: with low noise, nodes are not powerful enough to interact, producing a functional disconnection that was captured by the FC and dFC matrices. In sharp contrast, in postbifurcation, nodes were self-oscillating in alpha frequency around 10Hz. Note the similarity between high and low noise to the thalamus in postbifurcation range, supporting the results reported in previous sections.

### Thalamic alpha propagates to the cortex within a balanced SNR

The best-performing thalamocortical model was the one in prebifurcation that integrated high thalamic noise and pTh. Looking at its underlying dynamics, we observed that the spectrum was showing a 1/f shape. As we are trying to reproduce MEG FC, in which a predominant alpha frequency is usually observed, we wondered whether a spectral change towards alpha would impact the model performance. In this section, we manipulated the oscillatory frequency of the thalamus making it surpass bifurcation and self-oscillate in alpha by varying its average input, p_th_. These simulations were performed with pTh structure and *η_th_* = 0.022.

Simulations rising p_th_ in prebifurcation (g<7), resulted in a transition of thalamic nodes from the noisy 1/f pink noise spectra to an alpha oscillation (see Fig. 5, FFT peak), passing first through the slow and high amplitude limit cycle of the JR model [55, 56] at p_th_=[0.11 - 0.13] (see the dark orange spot in Fig. 5 SNR). In this transition along the bifurcation, r_PLV_ lowered down right after the high power and slow limit cycle (p_th_ > 0.13). This could be due to a high SNR ≈ 6 producing hypersynchronization (mean PLV≈0.7; see Fig. 5 mean PLV). However, we did not find this phenomenon with the slow limit cycle (PLV mean ≈0.5) even though it showed a higher SNR. This might be explained by the alpha band filtering that leaves out of the analysis the potentially hypersynchronizing oscillations of the slow limit cycle.

**Fig 5.**
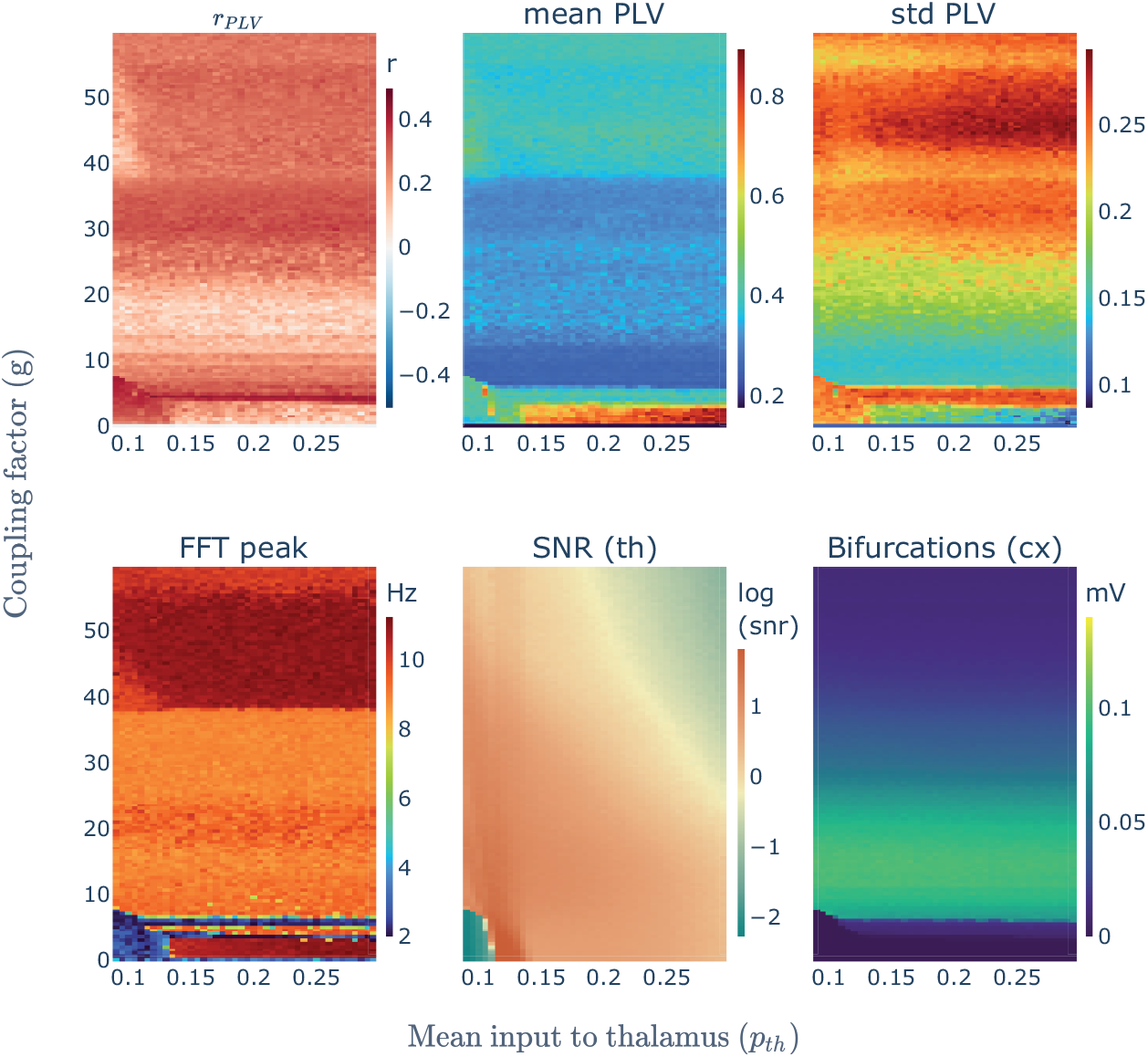
Parameter space explorations for p_th_ with the high thalamic noise model. Sets of three simulations averaged for subject one and parceled thalamus. Heatmaps show several metrics characterizing each simulation in the parameter space.

Interestingly, the changes in thalamic activity indirectly increased the inter-regional inputs to cortical nodes, making some of them pass bifurcation at g≈3 and g≈6 (see the horizontal blue lines in Fig. 5, FFT peak). These nodes produced a further rise in r_PLV_ (see the horizontal red line in Fig. 5, r_PLV_) suggesting two important things for prebifurcation simulations: 1) that every node is a potential contributor of a general driving mechanism that we have located in the thalamus (through a high noise), and 2) that the number of drivers participating in that mechanism matters.

In the range of p_th_ = [0.13-0.3], where we observed alpha oscillations, r_PLV_ decreased. We wondered whether this effect could be related to the rise in SNR after setting the thalamus to oscillate in the alpha band. Therefore, we fixed p_th_=0.15, and we varied the noisy input to the thalamus (*η_h_*) to explore its impact on model performance. We found an optimal balance for SNR at η_th_=[0.05-0.15] in which the noisy inputs to the thalamus were enough to avoid hypersynchronization (see Fig. 6 mean PLV) and low enough to maintain the intrinsic dynamics produced by the thalamus (i.e., alpha oscillations, see Fig. 6 FFT peak). Further increases in noise would replace progressively the alpha oscillatory behavior by a 1/f spectra without reducing r_PLV_.

**Fig 6.**
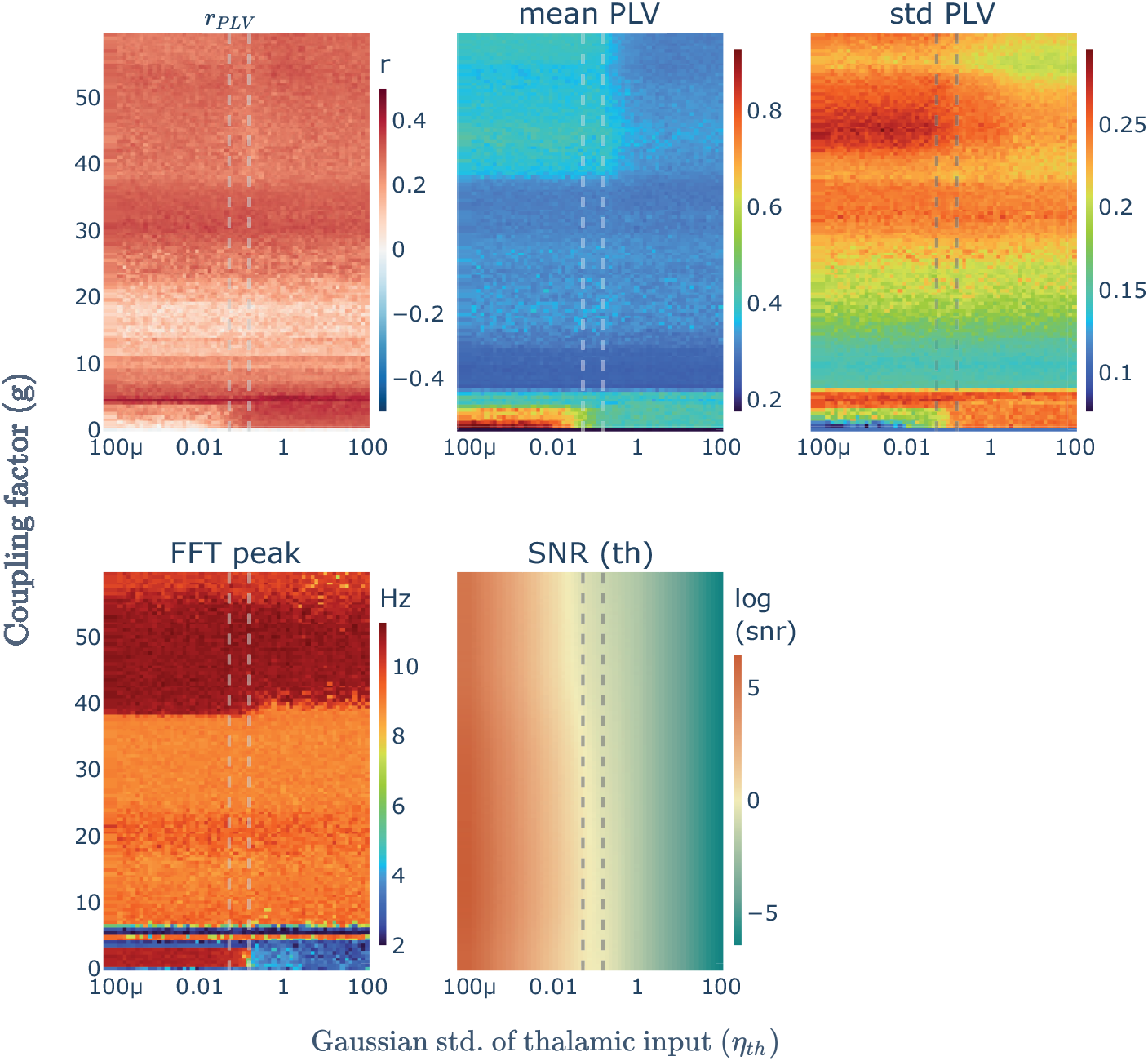
Parameter space explorations over *η_th_* to balance SNR in the thalamus. Sets of three simulations averaged for subject one and parceled thalamus. Heatmaps show several metrics characterizing each simulation in the parameter space. Vertical dashed lines define the optimal SNR range.

From these observations, it could be thought that a general rise of noise in the model (i.e., to all nodes) would lead to a better performance, however, in our modeling framework this is only true when the noise is implemented into a limited number of nodes. Independent noise into all cortical nodes is not linked to an enhancement of r_PLV_ (see supplementary Figure S3 Fig).

From these parameter explorations, we extracted four additional models of interest to explore their underlying dynamics (see Fig.7). Two of them related to the exploration of p_th_: one for the hypersynchrony situation (p_th_=0.15, *η_th_*=0.022), another for the slow JR limit cycle (p_th_=0.12, *η_th_* =0.022); and another two regarding the exploration of SNR: one for the optimal SNR (p_th_=0.15, *η_th_*=0.09), and another for a higher noise than the optimal range (p_th_=0.15, *η_th_*=0.5). Figure 7 shows the underlying dynamics for each of the additional models scenarios simulated with g=2.

**Fig 7.**
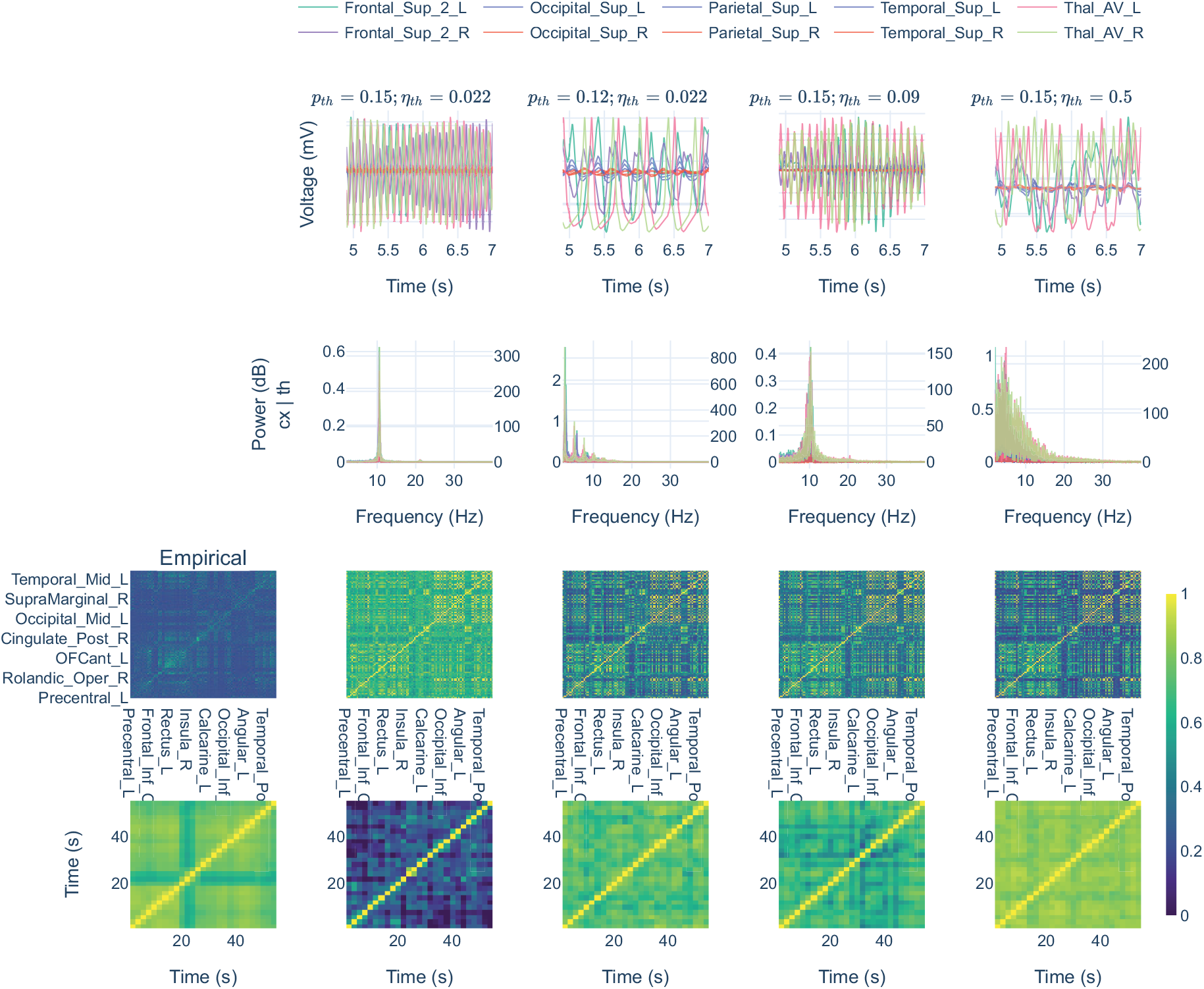
Complementary simulation samples. Simulation samples derived from the parameter explorations show additional situations in which the thalamus drives the cortical activity. Simulations for one subject, g = 2 and pTh. Rows showing demeaned signals, spectra, PLV, and dFC matrices, respectively.

## Discussion

Understanding thalamocortical networks is crucial for unraveling the complex dynamics of the human brain. These networks are essential for transmitting sensory information from the periphery to the cortex and regulating cortical arousal, which are fundamental processes for perception, attention, and cognition. In this study, we aimed to gain a deeper understanding of the role of the thalamus in rsFC by utilizing computational brain models. Our experiments focused on testing two key features of thalamocortical networks: the presence of thalamic afferences related to the RAS and first-order sensory relays, and the structure of the thalamus. Interestingly, we found that only when we raised thalamic noisy inputs to represent the RAS and peripheral afferences activating a damped cortex, its presence affected the simulated dynamics. To validate our findings, we performed a control experiment using the cerebro-cerebellar network and showed that implementing high noise into the cerebellum could replicate the results observed in the thalamocortical network. Finally, we explored how the oscillatory behavior of thalamic nodes, specifically their frequency and SNR, could shape the emergent rsFC. Our study provides novel insights into the role of thalamocortical networks in shaping brain dynamics and highlights the relevance of balanced SNR activity for the propagation of alpha rhythms from the thalamus.

We expected that introducing the thalamus in our simulations would have generated a difference in model performance in postbifurcation, where the best correlations are usually found. However, this was not the case, as implementing the thalamus in our model only affected results in prebifurcation and when introducing a high thalamic noise to represent afferences from RAS system and peripheral sensory relays. In prebifurcation, nodes are operating as damped oscillators, tending to lower down their activity and oscillation amplitude to a fixed point. If we introduce high noise in the thalamus, thalamic afferences would rise the activity in the cortex and allow for functional interactions. In other words, the thalamus would be driving cortical activation. The best model performance in this study was observed within this parametrization and including the thalamus with its dorsal nuclei divisions (pTh).

A further control experiment demonstrated that this mechanism was not directly related to thalamic SC. This was ascertained by embedding the driver mechanism into another hub-like region (i.e., cerebellum) and obtaining equivalent results for the match to empirical rsFC. This would mean that any node could theoretically play the role that we are attributing to the thalamus. In this line, during parameter explorations, some isolated cortical nodes passing bifurcation would contribute to enhance performance by being part of that driving mechanism. This would add up to the observation that parceled structures (i.e., thalamus and cerebellum) performed better than their single node versions, indicating that the number of driver nodes matters. In further research, it should be explored whether there is a computational optimum number of drivers to reproduce rsFC that could be subject or session-specific, and whether these differences may be linked to differences in brain and cognitive functioning.

Taken together, our main conclusion is that *a limited set of driving nodes is likely to underlie the dynamics of rsFC.* We believe that those drivers might be linked both to the regions participating in the RAS system (Intralaminar Thalamus, Raphe Nuclei, and Locus Coeruleus) [4, 16] and to the dorsal nuclei of the thalamus that are implicated in the relay of sensory information and have also been tightly linked to the generation of oscillatory behavior in the cortex in slow waves [63–66]. This would support the view of the thalamus as a driver and controller of cortical dynamics [6, 25, 67–69].

Further explorations on the spectral characteristics of the drivers showed that the thalamus could propagate its own intrinsic alpha dynamics when a balanced SNR was achieved. This feature represents the interaction between thalamocortical pacemaker neurons [14] and peripheral sensory inputs reaching the system and provoking event-related desynchronization [70, 71]. The model showed a good performance for reproducing empirical rsFC in that optimal range and with additional noise, generating a 1/f spectra. However, lower levels of noise with an alpha-oscillating thalamus reduced performance and led to a hypersynchronization situation transmitted from the thalamus to the cortex, generating an epileptic-like dynamic [72, 73]. In line with these results, the thalamus has been proposed to be involved in the onset of temporal lobe epileptic seizures, transmitting more regular patterns of activity to the hippocampus [74, 75].

From the computational perspective, previous work has shown that BNMs may reproduce better empirical rsFC when the models operate at the edge of bifurcation [59], the critical point. At this point, noisy excursions or the effect of nodes’ interaction can lead the masses to behave in any of the two regimes separated by the critical point. This phenomenon, referred to as criticality [58], has been proposed to enhance the capacity of brain systems to convey information [76]. However, in our study, we showed an equivalent performance both at the edge of bifurcation and over the whole prebifurcation range. This contrast might be explained by the different cortical and thalamic parametrization implemented in our nodes. Our thalamic driving nodes feed the cortex, leading the dynamics. In the cited study [59], the nodes that randomly switch between states would represent the same driving mechanism as in our model. Interestingly though, our approach would suggest that criticality is not a necessary feature in the dynamics of a resting brain at the mesoscale level.

Many studies in the field have explicitly [78–80] or implicitly [58, 81–83] excluded subcortical regions from their BNMs. This could be due to the technical limitations of recording deep brain signals and/or to the complexity of reconstructing SC schemes for small, deep crossing-fibers regions. But, more importantly, it could be due to the unknown role that these regions may play in shaping simulated whole brain dynamics. Some studies have attempted to unravel these mechanisms, showing the importance of the cerebellum for brain dynamics [60], the relevance of cortico-subcortical interaction for shaping dynamical functional connectivity [84] and the relevance of neurotransmission [57, 85]. Here, we described the potential role of the thalamus (and other activating brain regions) in whole brain simulations, linked to both neurotransmission and relay of information.

In conclusion, our study provides novel insights into the role of thalamocortical networks in shaping brain dynamics. We demonstrate that a limited set of driving nodes leading cortical activation may better describe resting-state activity. The thalamus would be a relevant part in this mechanism due to its participation in the RAS system and through its peripheral sensory relays being delivered from its multiple dorsal nuclei. In this type of architecture, driving nodes might show a balanced SNR to avoid hypersynchronization in the network. Although it is still debated whether the thalamus has an active role in cognition [6, 24, 86], our study strongly suggests its active participation in driving cortical dynamics and shaping FC in resting-state. These findings may contribute to a better understanding of brain function and dysfunction, fostering the development of new therapeutic approaches targeting thalamocortical circuits.

## Materials and methods

### Empirical dataset

MRI (T1 and DWI) scans and MEG recordings were acquired from 10 healthy participants in resting-state, with ages between 62 and 77 years old (mean 69, sd 4.17, 3 males, 7 females) from a dataset owned by the Centre of Cognitive and Computational Neuroscience, UCM, Madrid.

MRI-T1 scans were recorded in a General Electric 1.5 Tesla magnetic resonance scanner, using a high-resolution antenna and a homogenization PURE filter (fast spoiled gradient echo sequence, with parameters: repetition time/echo time/inversion time=11.2/4.2/450 ms; flip angle=12°; slice thickness=1 mm, 256×256 matrix, and field of view=256 mm).

Diffusion-weighted images (dw-MRI) were acquired with a single-shot echo-planar imaging sequence with the parameters: echo time/repetition time=96.1/12,000 ms; NEX 3 for increasing the SNR; slice thickness=2.4 mm, 128× 128 matrix, and field of view=30.7 cm yielding an isotropic voxel of 2.4 mm; 1 image with no diffusion sensitization (i.e., T2-weighted b0 images) and 25 dw-MRI (b=900 s/mm2).

MEG recordings were acquired with an Elekta-Neuromag MEG system with 306 channels at 1000Hz sampling frequency and an online band-pass filtered between 0.1 and 330Hz. MEG protocol consisted of 5 min resting-state eyes closed.

All participants provided informed consent.

### Functional connectivity

MEG recordings were preprocessed offline using the spatiotemporal signal space separation (tSSS) filtering algorithm [87], embedded in the Maxfilter Software v2.2 (correlation limit of 0.9 and correlation window of 10 seconds), to eliminate magnetic noise and compensate for head movements during the recording. Continuous MEG data were preprocessed using the Fieldtrip Toolbox [88], where an independent component-based algorithm was applied to remove the effects of ocular and cardiac signals from the data, together with external noise.

Source reconstruction was performed using the software Brainstorm [89], anatomically informed by the MRI scans of each subject. We employed the minimum norm estimates method [90], with the *constrained dipoles* variant, by which the current dipoles are oriented normally to the cortical surface, to model the orientation of the macrocolumns of pyramidal neurons, perpendicular to the cortex [91]

Source-space signals were then filtered in the classical frequency bands: delta (2-4 Hz), theta (4-8 Hz), alpha (8-12 Hz), beta (12-30 Hz), and gamma (30-45 Hz). For each subject and frequency band, FC was calculated between the source time series using the Phase Locking Value (PLV, [92]), and the resulting matrices were averaged into the AAL2 parcellation scheme [93]. In addition, we computed dynamical functional connectivity matrices by extracting PLV on consecutive intervals of 4 seconds of length (using the sliding window approach with 50% of overlapping) and evaluating the correlation between these PLV matrices.

### Structural connectivity

Diffusion-weighted images were processed using DSI Studio (http://dsi-studio.labsolver.org). The quality of the images was checked before fiber tracking and corrected for motion artifacts, eddy currents, and phase distortions. Then, tensor metrics were calculated. To improve reproducibility, we used a deterministic fiber tracking algorithm with augmented tracking strategies [94–96]. The whole brain volume was used as seeding region. Both the anisotropy and angular thresholds were randomly selected (the latter, from 15 degrees to 90 degrees). The step size was randomly selected from 0.5 voxels to 1.5 voxels. A total of 5 million seeds were placed and tracks with lengths shorter than 15 or longer than 180 mm were discarded.

To explore the impact of including/excluding the thalamus in simulations, we performed a first experiment comparing three different structural connectivity (SC) versions of each subject brains’: woTh, Th, and pTh. pTh consists of a brain network with 148 regions extracted from AAL3 atlas from which we kept the thalamic parcellation and, removed and merged the other areas to make it comparable to AAL2 scheme; Th consists of the 120 regions from AAL2; same for woTh in which we removed thalamic nuclei (118 regions). These three versions of the structural connectomes are represented in figure 8, and a list with all ROIs included can be found in S1 Table. Two connectivity matrices were calculated per SC version: counting the number of tracts connecting (i.e., passing through) each pair of brain regions, and the average length of those tracts.

**Fig 8.**
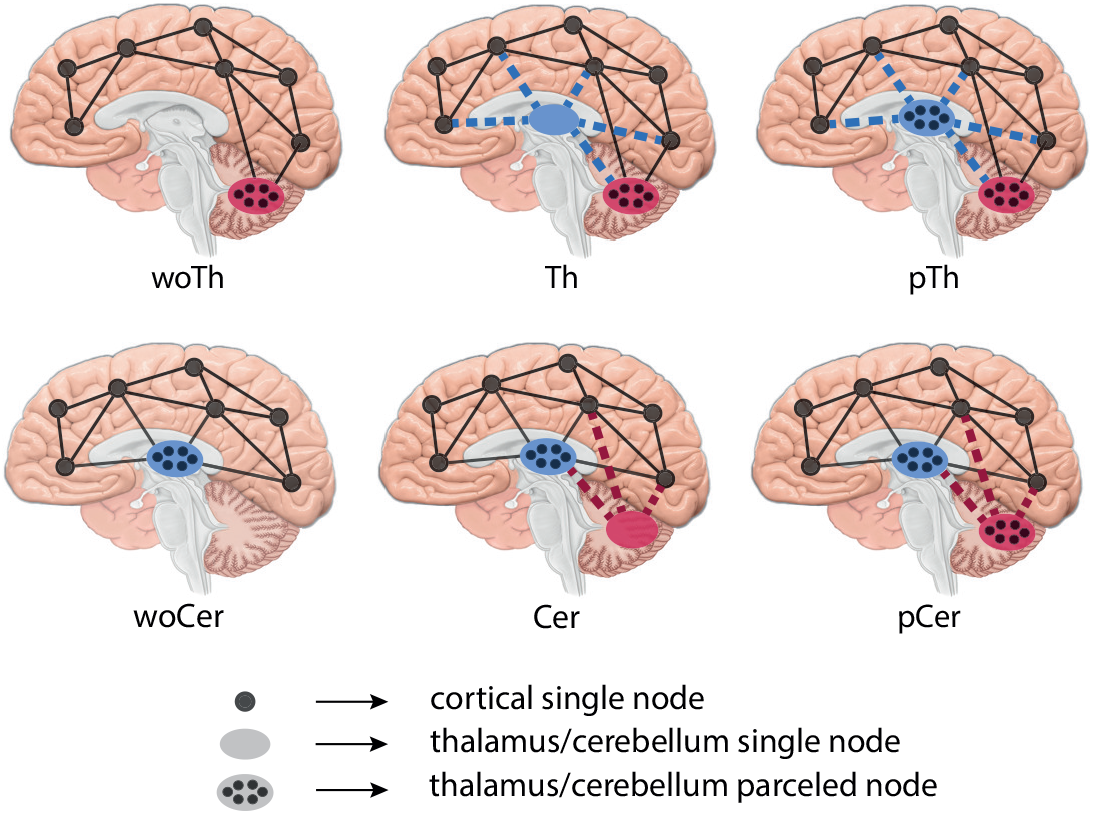
SC versions. First line shows the SC versions for the thalamocortical experiment: woTh, Th, pTh. The second line shows the SC versions for the cortico-cerebellar control experiment. Dashed lines representing driver connections.

As a control experiment, we applied the same process to the cerebellum as it is a brain region with similar network characteristics (see Table 1 and S1 Table), and it can also be modeled as a parceled structure and a single node. We extracted three SC versions per brain using AAL3 atlas: parceled cerebellum (pCer), cerebellum as a single node (Cer), and without cerebellum (woCer). The thalamus was modeled as parceled through all these versions, and therefore the resulting SCs were composed of 148, 122 and 120 regions, respectively.

### Brain network model

SC matrices served as the skeleton for the BNMs implemented in TVB [97] where regional signals were simulated using JR NMMs [54]. This is a biologically inspired model of a cortical column capable of reproducing alpha oscillations through a system of second-order coupled differential equations:

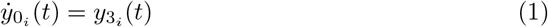

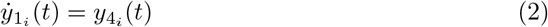

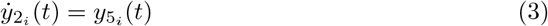

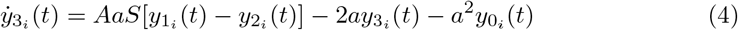

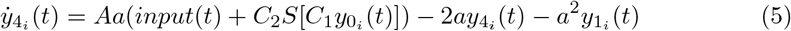

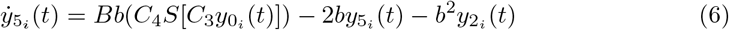

Where:

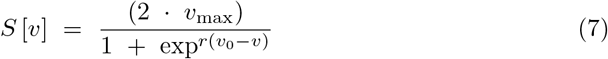

The inter-regional communication introduces heterogeneity in terms of connection strength *w_ji_*, and conduction delays *d_ji_* between nodes i and j, where:

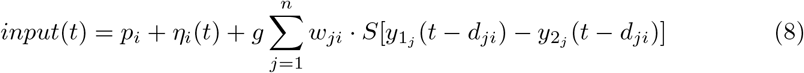

It represents the electrophysiological activity (in voltage) from three subpopulations of neurons: pyramidal neurons (*y*_0_), excitatory interneurons (*y*_1_), and inhibitory interneurons (*y*_2_). These subpopulations are interconnected (Fig 9) and integrate external inputs from other cortical columns. The communication is implemented in terms of firing rate (Eqs. 1 to 6) and a sigmoidal function (Eq. 7) stands for the conversion from voltage to firing rate.

**Fig 9.**
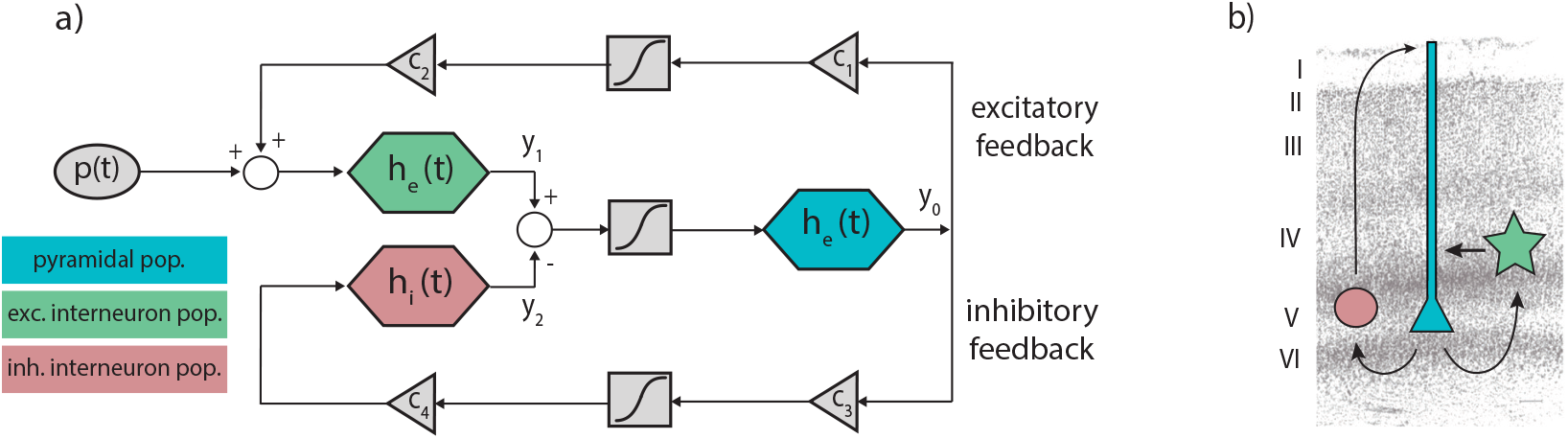
JR model of a cortical column. a) Block diagram depicting JR operators and modules where each color is associated with a different neural population: pyramidal (cyan), excitatory interneurons (green) and inhibitory interneuron (red). b) Histological contextualization of the cortical layers. Modified from [54, 81]

The input represents two main drivers of activity in the NMMs: inter-regional communication and intrinsic input. The latter is defined by a Gaussian noise with *p* mean and *η* std.

A global coupling factor *g* is implemented to scale linearly tracts’ weights. Both, *g* and *v* are scaling factors that apply to all nodes. Parameter values are described in table 2.

**Table 2.** JR parameters used in simulations.

| Parameter | Value | Unit | Description |
| --- | --- | --- | --- |
| A | 3.25 | mV | Average excitatory synaptic gain |
| B | 22 | mV | Average inhibitory synaptic gain |
| a | 0.1 | $\text{ms}^{-1}$ | Time Constant of excitatory PSP |
| b | 0.05 | $\text{ms}^{-1}$ | Time Constant of inhibitory PSP |
| C1 | 135 |  | Average synaptic contacts: pyramidal to excitatory interneurons |
| C2 | 108 |  | Average synaptic contacts: excitatory interneurons to pyramidal |
| C3 | 33.75 |  | Average synaptic contacts: pyramidal to inhibitory interneurons |
| C4 | 33.75 |  | Average synaptic contacts: inhibitory interneurons to pyramidal |
| v_max | 0.0 | $\text{ms}^{-1}$ | Half the maximum firing rate |
| r | 0.56 | $\text{mV}^{-1}$ | Slope of the presynaptic function at $v_0$ |
| $v_0$ | 6 | mV | Potential when half the maximum firing rate is achieved |
| p | variable | $\text{ms}^{-1}$ | Mean of random Gaussian intrinsic input |
| $\eta$ | variable | $\text{ms}^{-1}$ | Standard deviation of random Gaussian intrinsic input (Noise) |
| g | variable |  | Coupling factor for inter-regional communication - multiplier of weights- |
| s | 15 | mm/ms | Conduction speed for inter-regional communication |

Unless otherwise stated, we used default values for parameters $p = 0.09$ and $\eta = 2.2 \times 10^{-8}$ . Note that along the study, we introduce bimodalities in those parameters $p_{th}$ and $\eta_{th}$ )

The JR model shows two supercritical hopf bifurcations for the parameter *p* [56]. When JR NMMs are implemented in a connected network, the parameter *g* scales the inter-regional input to nodes, becoming a bifurcation parameter. We used the first bifurcation to separate two NMM’s behaviors (Fig. 10): damped oscillator in the *prebifurcation* range where nodes tend to a fixed point in voltage, and limit cycle in the *postbifurcation* range where nodes self-oscillate in the alpha frequency.

**Fig 10.**
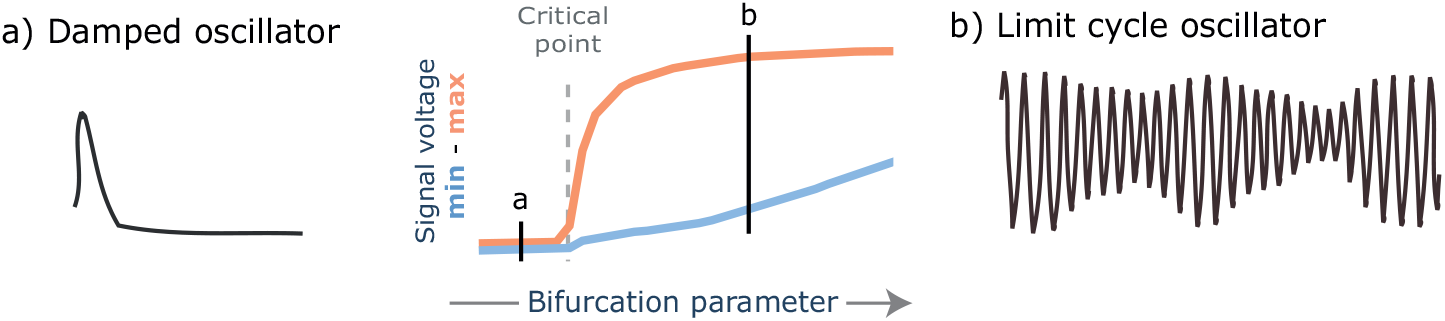
The bifurcation separates two states. In the center, a bifurcation diagram shows the minimum and maximum voltage for each value of the bifurcation parameter. At the critical point (dashed line), the bifurcation occurs and separates two states of the system: a) damped oscillator, whose activity tends to decay to a fixed point; and b) limit cycle oscillator, whose activity is a self-sustained oscillation.

### Simulations

In the first experiment, simulations were performed varying two parameters: the standard deviation of the input to the thalamus (i.e., noise, *η_th_*=[0.022, 2.2×10^-8^]) representing the presence/absence of subcortical and peripheral inputs; and the thalamic structure (woTh, Th, pTh). Additionally, we explored the parameter space for coupling factor (g=[0-60]). These models were simulated with the parameter p set to 0.09. We performed three simulations of 60 seconds (removing the initial 4 seconds to avoid transients) per model and computed two metrics: the Pearson’s correlation coefficient between the vectorized upper triangular matrices of the simulated and empirical FC (i.e., r_PLV_); and the KSD between the distributions of correlations in the dFC matrices (empirical and simulated). We restricted the PLVs’ comparisons to 1) only cortical regions, to avoid the limitations of MEG recordings regarding deep brain signals; and 2) only the alpha band, for computational simplicity and to be aware that it dominates MEG resting-state recordings. The same configuration was used in the control experiment with the cerebellum.

For further explorations, we simulated for 10 seconds (omitting initial 2 seconds to avoid transients) different ranges of the mean intrinsic input to the thalamus p_th_) and its standard deviation (*η_th_*). Besides the rPLV and KSD metrics, we show 1) *bifurcation diagrams* capturing the averaged maximum and minimum signal’s voltage per simulated ROI at each point in the parameter space. In the case of exploring 2 parameters at the same time (e.g., *g* and p_th_), bifurcations are presented in heatmaps conveying information about the difference between the maximum and minimum signal voltage; 2) *signal-to-noise ratio* (SNR) in the thalamus that is computed by dividing the amplitude of simulated signals by the standard deviation of the Gaussian noise used for the thalamus; 3) *relative power* between cortex and thalamus calculated by dividing the averaged area of cortical spectra by the averaged area of thalamic spectra.

### Statistics

We averaged the results of the 3 sets of simulations (i.e., repetitions) and performed statistical analysis for the group of 10 subjects. For the first experiment, we considered the maximum r_PLV_ and minimum KSD per subject, thalamic SC version and scenario. The effects of thalamic SC version and noise levels were evaluated using four two-way repeated measures ANOVA: two comparing maximum r_PLV_ in prebifurcation and postbifurcation; and another two comparing minimum KSD; after checking for the statistical assumptions of normality (Shapiro’s test) and sphericity (Mauchly’s test). Pairwise comparisons for thalamic structure and thalamic noise were evaluated using Wilcoxon test, correcting for multiple comparisons using FDR Benjamini-Hochberg method. The same procedure was applied in the control experiment with the cerebro-cerebellar network to maximum r_PLV_ comparisons.

## Acknowledgments

Thanks to Claudio Mirasso for his useful comments on the text. This research was funded by the Spanish Ministry of Universities through a predoctoral FPU grant to JCA (FPU2019-04251).

## Supporting information

**S1 Fig.**
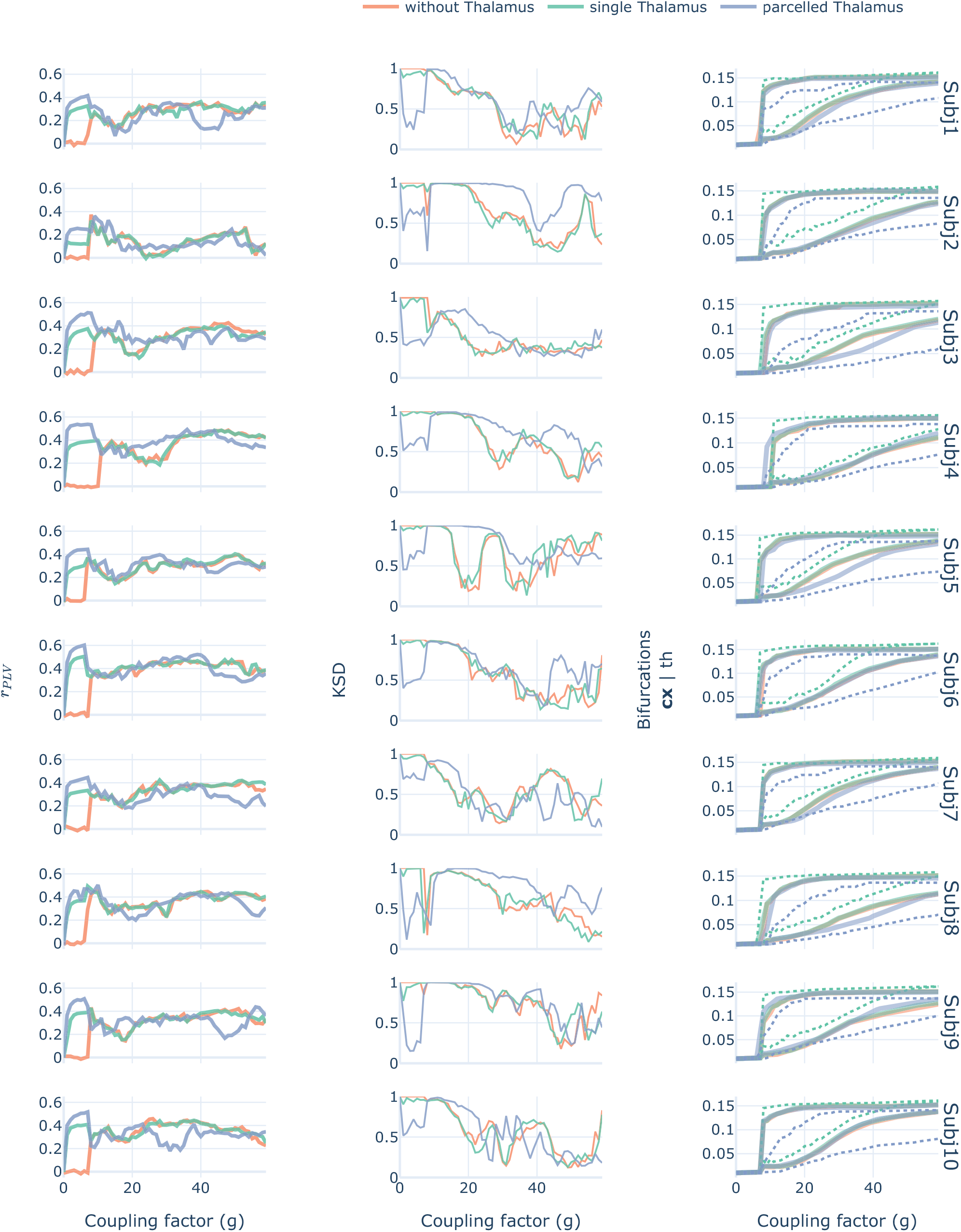
Lineplots showing r_PLV_, KSD and bifurcations per subject and SC version. The global behavior in r_PLV_ (first column) was similar for every subject. Note slight differences for subject 2 and subject 8 in which the bifurcation does not match the highest r_PLV_ value.

**S2 Fig.**
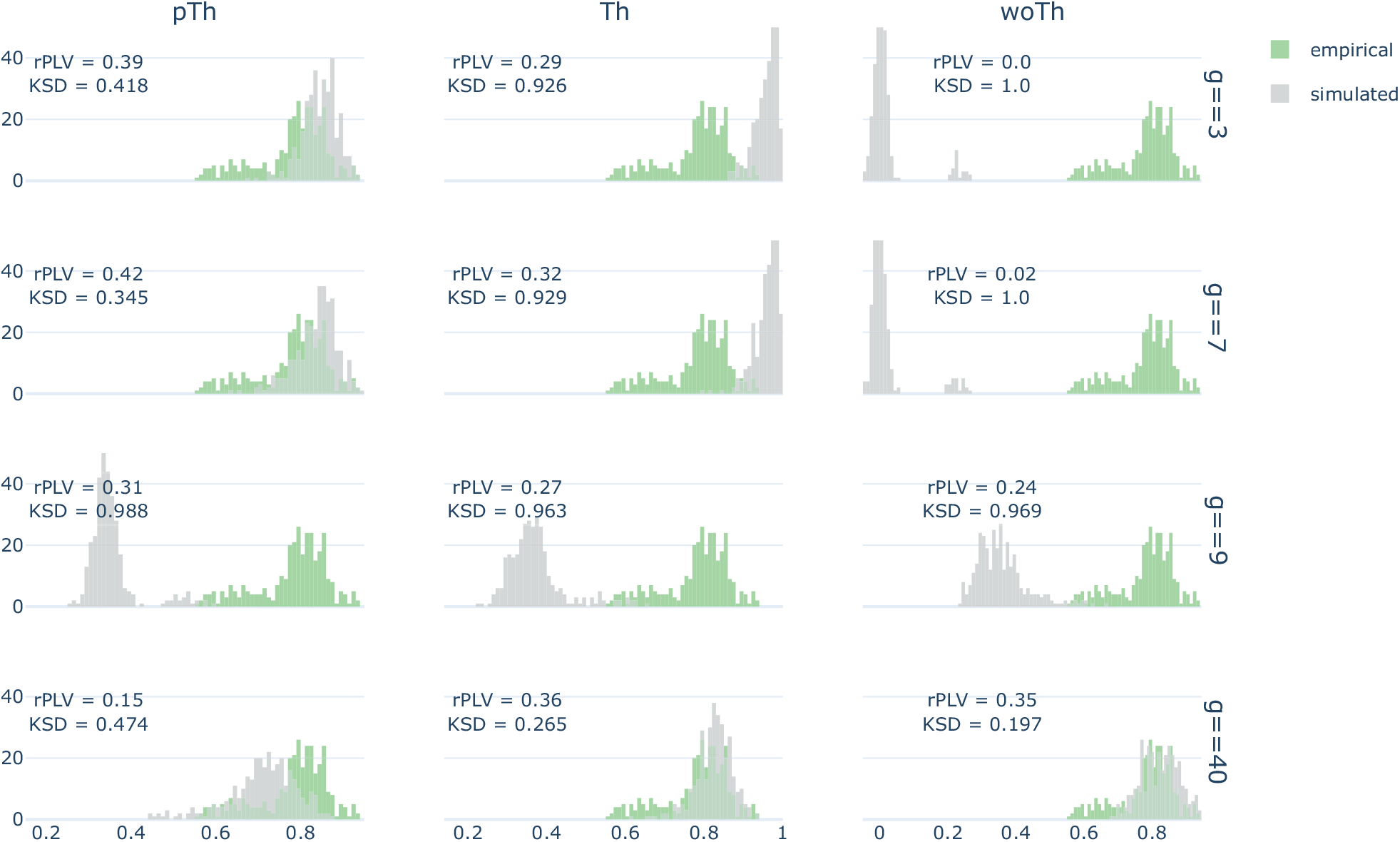
Empirical and simulated distributions of Pearson’s correlation values in dFC matrices. Simulations were performed for subject 1 with high thalamic noise (*η_th_* =0.022) and in both prebifurcation (*g*=[3, 7]) and postbifurcation 2 (*g*=[9, 40]). The three thalamocortical SC (i.e., woTh, Th, pTh) versions were simulated.

**S3 Fig.**
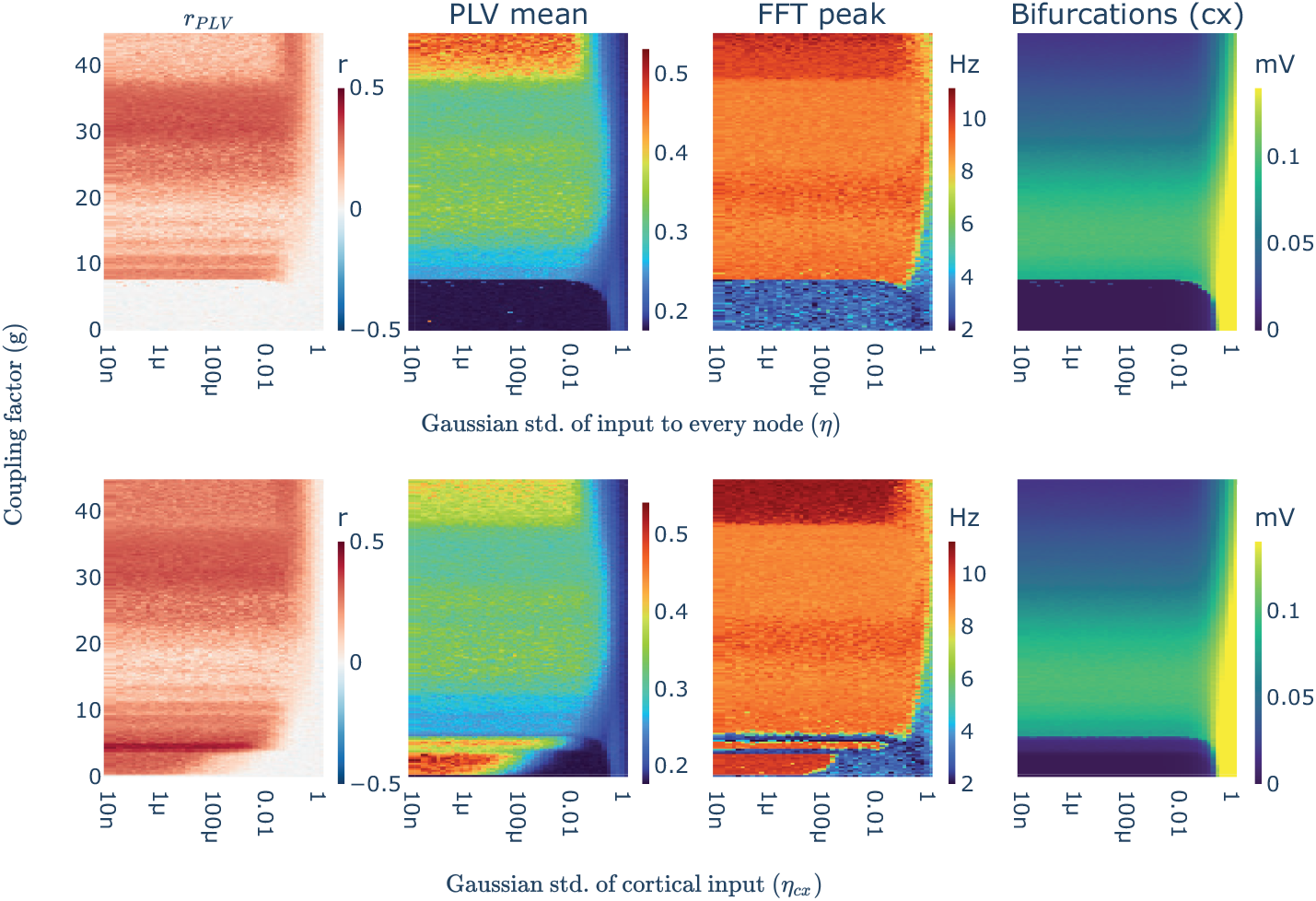
Rising cortical noise hampers r_PLV_. First row, showing simulations where all nodes have the same parametrization (*p*=0.09; *η*=variable). Second row, showing simulations with the thalamus in limit cycle condition p_th_=0.15, *η_th_*=0.09) and a variable noisy input to cortical nodes (*p_cx_*=0.09, *η_cx_*=variable).

**S1 Table. Network analysis of the regions included in the BNMs.** Degree, the number of neighbors of a region, and node strength, the average number of streamlines connecting a region to others were normalized over their respective maxima. Betweenness captures the number of shortest paths in a network that passes through a node. Path length stands for the average of the shortest paths for a node. Metrics were calculated with Networkx package [98] in Python 3.9. The thalamus and the cerebellum are considered here in the parcellated version. Note that Cingulate_Ant in AAL3 is divided in 3 parts and it was merged to match AAL2 scheme.

## References

1. Guillery RW, Sherman SM. Thalamic Relay Functions and Their Role in Corticocortical Communication. Neuron. 2002;33(2):163–175. doi:10.1016/s0896-6273(01)00582-7.

2. Sherman SM, Guillery RW. Distinct functions for direct and transthalamic corticocortical connections. Journal of Neurophysiology. 2011;106(3):1068–1077. doi:10.1152/jn.00429.2011.

3. Kato N. Cortico-thalamo-cortical projection between visual cortices. Brain Res. 1990;509:150–2.

4. Garcia-Rill E. Reticular Activating System. In: Encyclopedia of Neuroscience. Elsevier; 2009. p. 137–143. Available from: https://doi.org/10.1016%2Fb978-008045046-9.01767-8.

5. Jones EG. Principles of Thalamic Organization. In: The Thalamus. Springer US; 1985. p. 85–149. Available from: https://doi.org/10.1007%2F978-1-4615-1749-8_3.

6. Sherman S. Thalamus plays a central role in ongoing cortical functioning. Nat Neurosci. 2016;19:533–41.

7. Halassa M, Sherman S. Thalamocortical Circuit Motifs: A General Framework. Neuron. 2019;103:762–770.

8. Crabtree JW. Functional Diversity of Thalamic Reticular Subnetworks. Frontiers in Systems Neuroscience. 2018;12. doi:10.3389/fnsys.2018.00041.

9. Sherman SM, Guillery RW. Exploring the Thalamus and Its Role in Cortical Function. Second edition ed. MIT press; 2005.

10. Schreiner T, Kaufmann E, Noachtar S, Mehrkens JH, Staudigl T. The human thalamus orchestrates neocortical oscillations during NREM sleep. Nature Communications. 2022;13(1). doi:10.1038/s41467-022-32840-w.

11. Lopes da Silva F. Neural mechanisms underlying brain waves: from neural membranes to networks. Electroencephalogr Clin Neurophysiol. 1991;79:81–93.

12. David O, Friston K. A neural mass model for MEG/EEG: coupling and neuronal dynamics. Neuroimage. 2003;20:1743–55.

13. Fuentealba P, Steriade M. The reticular nucleus revisited: Intrinsic and network properties of a thalamic pacemaker. Progress in Neurobiology. 2005;75(2):125–141. doi:10.1016/j.pneurobio.2005.01.002.

14. Fogerson PM, Huguenard JR. Tapping the Brakes: Cellular and Synaptic Mechanisms that Regulate Thalamic Oscillations. Neuron. 2016;92(4):687–704. doi:10.1016/j.neuron.2016.10.024.

15. Huguenard J, McCormick D. Thalamic synchrony and dynamic regulation of global forebrain oscillations. Trends Neurosci. 2007;30:350–6.

16. Steriade M. Arousal–Revisiting the Reticular Activating System. Science. 1996;272(5259):225–225. doi:10.1126/science.272.5259.225.

17. Iturria-Medina Y, Canales-Rodríguez E, Melie-García L, Valdés-Hernández P, Martínez-Montes E, Alemán-Gómez Y, et al. Characterizing brain anatomical connections using diffusion weighted MRI and graph theory. Neuroimage. 2007;36:645–60.

18. Behrens T, Johansen-Berg H, Woolrich M, Smith S, Wheeler-Kingshott C, Boulby P, et al. Non-invasive mapping of connections between human thalamus and cortex using diffusion imaging. Nat Neurosci. 2003;6:750–7.

19. Hwang K, Bertolero MA, Liu WB, D’Esposito M. The Human Thalamus Is an Integrative Hub for Functional Brain Networks. The Journal of Neuroscience. 2017;37(23):5594–5607. doi:10.1523/jneurosci.0067-17.2017.

20. Vantomme G, Osorio-Forero A, Lüthi A, Fernandez L. Regulation of Local Sleep by the Thalamic Reticular Nucleus. Front Neurosci. 2019;13:576.

21. Jan J, Reiter R, Wasdell M, Bax M. The role of the thalamus in sleep, pineal melatonin production, and circadian rhythm sleep disorders. J Pineal Res. 2009;46:1–7.

22. Jin Y, Yang H, Zhang F, Wang J, Liu H, Yang X, et al. The Medial Thalamus Plays an Important Role in the Cognitive and Emotional Modulation of Orofacial Pain: A Functional Magnetic Resonance Imaging-Based Study. Front Neurol. 2020;11:589125.

23. Vartiainen N, Perchet C, Magnin M, Creac’h C, Convers P, Nighoghossian N, et al. Thalamic pain: anatomical and physiological indices of prediction. Brain. 2016;139:708–22.

24. Wolff M, Vann SD. The Cognitive Thalamus as a Gateway to Mental Representations. The Journal of Neuroscience. 2018;39(1):3–14. doi:10.1523/jneurosci.0479-18.2018.

25. Schmitt L, Wimmer R, Nakajima M, Happ M, Mofakham S, Halassa M. Thalamic amplification of cortical connectivity sustains attentional control. Nature. 2017;545:219–223.

26. Wimmer R, Schmitt L, Davidson T, Nakajima M, Deisseroth K, Halassa M. Thalamic control of sensory selection in divided attention. Nature. 2015;526:705–9.

27. Portas C, Rees G, Howseman A, Josephs O, Turner R, Frith C. A specific role for the thalamus in mediating the interaction of attention and arousal in humans. J Neurosci. 1998;18:8979–89.

28. Torrico T, Munakomi S. Neuroanatomy, Thalamus. In: StatPearls [Internet]. Treasure Island (FL): StatPearls Publishing; 2021. Available from: https://www.ncbi.nlm.nih.gov/books/NBK542184/.

29. Laureys S. The neural correlate of (un)awareness: lessons from the vegetative state. Trends Cogn Sci. 2005;9:556–9.

30. Alkire MT, Hudetz AG, Tononi G. Consciousness and Anesthesia. Science. 2008;322(5903):876–880. doi:10.1126/science.1149213.

31. Tasserie J, Uhrig L, Sitt J, Manasova D, Dupont M, Dehaene S, et al. Deep brain stimulation of the thalamus restores signatures of consciousness in a nonhuman primate model. Sci Adv. 2022;8:eabl5547.

32. Schiff N. Recovery of consciousness after brain injury: a mesocircuit hypothesis. Trends Neurosci. 2010;33:1–9.

33. Redinbaugh MJ, Phillips JM, Kambi NA, Mohanta S, Andryk S, Dooley GL, et al. Thalamus Modulates Consciousness via Layer-Specific Control of Cortex. Neuron. 2020;106(1):66–75.e12. doi:10.1016/j.neuron.2020.01.005.

34. Sherman SM. The thalamus is more than just a relay. Current Opinion in Neurobiology. 2007;17(4):417–422. doi:10.1016/j.conb.2007.07.003.

35. Nakajima M, Halassa MM. Thalamic control of functional cortical connectivity. Current Opinion in Neurobiology. 2017;44:127–131. doi:10.1016/j.conb.2017.04.001.

36. Barson JR, Mack NR, Gao WJ. The Paraventricular Nucleus of the Thalamus Is an Important Node in the Emotional Processing Network. Frontiers in Behavioral Neuroscience. 2020;14. doi:10.3389/fnbeh.2020.598469.

37. Dehghani N, Wimmer RD. A Computational Perspective of the Role of the Thalamus in Cognition. Neural Computation. 2019;31(7):1380–1418. doi:10.1162/neco_a_01197.

38. Friston KJ. Functional and effective connectivity in neuroimaging: A synthesis. Human Brain Mapping. 1994;2(1-2):56–78. doi:10.1002/hbm.460020107.

39. Friston KJ. Functional and Effective Connectivity: A Review. Brain Connectivity. 2011;1(1):13–36. doi:10.1089/brain.2011.0008.

40. Raichle ME. Two views of brain function. Trends in Cognitive Sciences. 2010;14(4):180–190. doi:10.1016/j.tics.2010.01.008.

41. Raichle ME, MacLeod AM, Snyder AZ, Powers WJ, Gusnard DA, Shulman GL. A default mode of brain function. Proceedings of the National Academy of Sciences. 2001;98(2):676–682. doi:10.1073/pnas.98.2.676.

42. Mason MF, Norton MI, Horn JDV, Wegner DM, Grafton ST, Macrae CN. Wandering Minds: The Default Network and Stimulus-Independent Thought. Science. 2007;315(5810):393–395. doi:10.1126/science.1131295.

43. Raichle ME. The Brain’s Default Mode Network. Annual Review of Neuroscience. 2015;38(1):433–447. doi:10.1146/annurev-neuro-071013-014030.

44. Christoff K, Gordon AM, Smallwood J, Smith R, Schooler JW. Experience sampling during fMRI reveals default network and executive system contributions to mind wandering. Proceedings of the National Academy of Sciences. 2009;106(21):8719–8724. doi:10.1073/pnas.0900234106.

45. Spinosa V, Brattico E, Campo F, Logroscino G. A systematic review on resting state functional connectivity in patients with neurodegenerative disease and hallucinations. NeuroImage: Clinical. 2022;35:103112. doi:10.1016/j.nicl.2022.103112.

46. Hausman HK, O’Shea A, Kraft JN, Boutzoukas EM, Evangelista ND, Etten EJV, et al. The Role of Resting-State Network Functional Connectivity in Cognitive Aging. Frontiers in Aging Neuroscience. 2020;12. doi:10.3389/fnagi.2020.00177.

47. Tang S, Wang Y, Liu Y, Chau SW, Chan JW, Chu WC, et al. Large-scale network dysfunction in *α*-Synucleinopathy: A meta-analysis of resting-state functional connectivity. eBioMedicine. 2022;77:103915. doi:10.1016/j.ebiom.2022.103915.

48. Betzel RF, Byrge L, He Y, Goñi J, Zuo XN, Sporns O. Changes in structural and functional connectivity among resting-state networks across the human lifespan. NeuroImage. 2014;102:345–357. doi:10.1016/j.neuroimage.2014.07.067.

49. Sala-Llonch R, Bartrés-Faz D, Junqué C. Reorganization of brain networks in aging: a review of functional connectivity studies. Frontiers in Psychology. 2015;6. doi:10.3389/fpsyg.2015.00663.

50. Fjell AM, Sneve MH, Grydeland H, Storsve AB, de Lange AMG, Amlien IK, et al. Functional connectivity change across multiple cortical networks relates to episodic memory changes in aging. Neurobiology of Aging. 2015;36(12):3255–3268. doi:10.1016/j.neurobiolaging.2015.08.020.

51. Fama R, Sullivan EV. Thalamic structures and associated cognitive functions: Relations with age and aging. Neuroscience & Biobehavioral Reviews. 2015;54:29–37. doi:10.1016/j.neubiorev.2015.03.008.

52. Goldstone A, Mayhew SD, Hale JR, Wilson RS, Bagshaw AP. Thalamic functional connectivity and its association with behavioral performance in older age. Brain and Behavior. 2018;8(4):e00943. doi:10.1002/brb3.943.

53. Schirner M, Rothmeier S, Jirsa V, McIntosh A, Ritter P. An automated pipeline for constructing personalized virtual brains from multimodal neuroimaging data. Neuroimage. 2015;117:343–57.

54. Jansen B, Rit V. Electroencephalogram and visual evoked potential generation in a mathematical model of coupled cortical columns. Biol Cybern. 1995;73:357–66.

55. Spiegler A, Kiebel S, Atay F, Knösche T. Bifurcation analysis of neural mass models: Impact of extrinsic inputs and dendritic time constants. Neuroimage. 2010;52:1041–58.

56. Grimbert F, Faugeras O. Bifurcation analysis of Jansen’s neural mass model. Neural Comput. 2006;18:3052–68.

57. Deco G, Cruzat J, Cabral J, Knudsen GM, Carhart-Harris RL, Whybrow PC, et al. Whole-Brain Multimodal Neuroimaging Model Using Serotonin Receptor Maps Explains Non-linear Functional Effects of LSD. Current Biology. 2018;28(19):3065–3074.e6. doi:10.1016/j.cub.2018.07.083.

58. Deco G, Jirsa V. Ongoing cortical activity at rest: criticality, multistability, and ghost attractors. J Neurosci. 2012;32:3366–75.

59. Deco G, Kringelbach M, Jirsa V, Ritter P. The dynamics of resting fluctuations in the brain: metastability and its dynamical cortical core. Sci Rep. 2017;7:3095.

60. Palesi F, Lorenzi R, Casellato C, Ritter P, Jirsa V, Gandini WKC, et al. The Importance of Cerebellar Connectivity on Simulated Brain Dynamics. Front Cell Neurosci. 2020;14:240.

61. Wendling F, Bartolomei F, Bellanger J, Chauvel P. Epileptic fast activity can be explained by a model of impaired GABAergic dendritic inhibition. Eur J Neurosci. 2002;15:1499–508.

62. Kiebel S, Garrido M, Moran R, Friston K. Dynamic causal modelling for EEG and MEG. Cogn Neurodyn. 2008;2:121–36.

63. Rigas P, Castro-Alamancos MA. Thalamocortical Up States: Differential Effects of Intrinsic and Extrinsic Cortical Inputs on Persistent Activity. Journal of Neuroscience. 2007;27(16):4261–4272. doi:10.1523/jneurosci.0003-07.2007.

64. Sanchez-Vives MV. Origin and dynamics of cortical slow oscillations. Current Opinion in Physiology. 2020;15:217–223. doi:https://doi.org/10.1016/j.cophys.2020.04.005.

65. Crunelli V, Hughes SW. The slow (<1 Hz) rhythm of non-REM sleep: a dialogue between three cardinal oscillators. Nat Neurosci. 2010;13(1):9–17.

66. Sheroziya M, Timofeev I. Global Intracellular Slow-Wave Dynamics of the Thalamocortical System. Journal of Neuroscience. 2014;34(26):8875–8893. doi:10.1523/jneurosci.4460-13.2014.

67. Mofakham S, Fry A, Adachi J, Stefancin PL, Duong TQ, Saadon JR, et al. Electrocorticography reveals thalamic control of cortical dynamics following traumatic brain injury. Communications Biology. 2021;4(1). doi:10.1038/s42003-021-02738-2.

68. Logiaco L, Abbott LF, Escola S. Thalamic control of cortical dynamics in a model of flexible motor sequencing. Cell Reports. 2021;35(9):109090. doi:10.1016/j.celrep.2021.109090.

69. Alonso J, Swadlow H. Thalamus controls recurrent cortical dynamics. Nat Neurosci. 2015;18:1703–4.

70. Pfurtscheller G, Neuper C, Mohl W. Event-related desynchronization (ERD) during visual processing. International Journal of Psychophysiology. 1994;16(2-3):147–153. doi:10.1016/0167-8760(89)90041-x.

71. Palva S, Palva JM. New vistas for *α*-frequency band oscillations. Trends in Neurosciences. 2007;30(4):150–158. doi:10.1016/j.tins.2007.02.001.

72. Jiruska P, de Curtis M, Jefferys JGR, Schevon CA, Schiff SJ, Schindler K. Synchronization and desynchronization in epilepsy: controversies and hypotheses. The Journal of Physiology. 2013;591(4):787–797. doi:10.1113/jphysiol.2012.239590.

73. Mishra AM, Bai X, Motelow JE, DeSalvo MN, Danielson N, Sanganahalli BG, et al. Increased resting functional connectivity in spike-wave epilepsy in WAG/Rij rats. Epilepsia. 2013;54(7):1214–1222. doi:10.1111/epi.12227.

74. Li YH, Li JJ, Lu QC, Gong HQ, Liang PJ, Zhang PM. Involvement of Thalamus in Initiation of Epileptic Seizures Induced by Pilocarpine in Mice. Neural Plasticity. 2014;2014:1–15. doi:10.1155/2014/675128.

75. Zhang CH, Sha Z, Mundahl J, Liu S, Lu Y, Henry TR, et al. Thalamocortical relationship in epileptic patients with generalized spike and wave discharges — A multimodal neuroimaging study. NeuroImage: Clinical. 2015;9:117–127. doi:10.1016/j.nicl.2015.07.014.

76. Cocchi L, Gollo L, Zalesky A, Breakspear M. Criticality in the brain: A synthesis of neurobiology, models and cognition. Prog Neurobiol. 2017;158:132–152.

77. Betzel RF, Bassett DS. Multi-scale brain networks. NeuroImage. 2017;160:73–83. doi:10.1016/j.neuroimage.2016.11.006.

78. David O, Harrison L, Friston K. Modelling event-related responses in the brain. Neuroimage. 2005;25:756–70.

79. Cabral J, Hugues E, Sporns O, Deco G. Role of local network oscillations in resting-state functional connectivity. Neuroimage. 2011;57:130–139.

80. Demirtaş M, Falcon C, Tucholka A, Gispert J, Molinuevo J, Deco G. A whole-brain computational modeling approach to explain the alterations in resting-state functional connectivity during progression of Alzheimer’s disease. Neuroimage Clin. 2017;16:343–354.

81. Stefanovski L, Triebkorn P, Spiegler A, Diaz-Cortes M, Solodkin A, Jirsa V, et al. Linking Molecular Pathways and Large-Scale Computational Modeling to Assess Candidate Disease Mechanisms and Pharmacodynamics in Alzheimer’s Disease. Front Comput Neurosci. 2019;13:54.

82. Courtiol J, Guye M, Bartolomei F, Petkoski S, Jirsa V. Dynamical Mechanisms of Interictal Resting-State Functional Connectivity in Epilepsy. J Neurosci. 2020;40:5572–5588.

83. Aerts H, Schirner M, Dhollander T, Jeurissen B, Achten E, Van RD, et al. Modeling brain dynamics after tumor resection using The Virtual Brain. Neuroimage. 2020;213:116738.

84. Favaretto C, Allegra M, Deco G, Metcalf N, Griffis J, Shulman G, et al. Subcortical-cortical dynamical states of the human brain and their breakdown in stroke. Nat Commun. 2022;13:5069.

85. Kringelbach ML, Cruzat J, Cabral J, Knudsen GM, Carhart-Harris R, Whybrow PC, et al. Dynamic coupling of whole-brain neuronal and neurotransmitter systems. Proceedings of the National Academy of Sciences. 2020;117(17):9566–9576. doi:10.1073/pnas.1921475117.

86. Sieveritz B, Raghavan R. The Central Thalamus: Gatekeeper or Processing Hub? J Neurosci. 2021;41:4954–4956.

87. Taulu S, Hari R. Removal of magnetoencephalographic artifacts with temporal signal-space separation: demonstration with single-trial auditory-evoked responses. Hum Brain Mapp. 2009;30:1524–34.

88. Oostenveld R, Fries P, Maris E, Schoffelen J. FieldTrip: Open source software for advanced analysis of MEG, EEG, and invasive electrophysiological data. Comput Intell Neurosci. 2011;2011:156869.

89. Tadel F, Baillet S, Mosher J, Pantazis D, Leahy R. Brainstorm: a user-friendly application for MEG/EEG analysis. Comput Intell Neurosci. 2011;2011:879716.

90. Hämäläinen M, Ilmoniemi R. Interpreting magnetic fields of the brain: minimum norm estimates. Med Biol Eng Comput. 1994;32:35–42.

91. Tadel F, Bock E, Niso G, Mosher J, Cousineau M, Pantazis D, et al. MEG/EEG Group Analysis With Brainstorm. Front Neurosci. 2019;13:76.

92. Lachaux J, Rodriguez E, Martinerie J, Varela F. Measuring phase synchrony in brain signals. Hum Brain Mapp. 1999;8:194–208.

93. Rolls E, Joliot M, Tzourio-Mazoyer N. Implementation of a new parcellation of the orbitofrontal cortex in the automated anatomical labeling atlas. Neuroimage. 2015;122:1–5.

94. Yeh F, Wedeen V, Tseng W. Generalized q-sampling imaging. IEEE Trans Med Imaging. 2010;29:1626–35.

95. Yeh FC, Verstynen TD, Wang Y, Fernández-Miranda JC, Tseng WYI. Deterministic Diffusion Fiber Tracking Improved by Quantitative Anisotropy. PLoS ONE. 2013;8(11):e80713. doi: 10.1371/journal.pone.0080713.

96. Yeh FC. Shape analysis of the human association pathways. NeuroImage. 2020;223:117329. doi:10.1016/j.neuroimage.2020.117329.

97. Sanz-Leon P, Knock S, Spiegler A, Jirsa V. Mathematical framework for large-scale brain network modeling in The Virtual Brain. Neuroimage. 2015;111:385–430.

98. Hagberg AA, Schult DA, Swart PJ. Exploring Network Structure, Dynamics, and Function using NetworkX. In: Varoquaux G, Vaught T, Millman J, editors. Proceedings of the 7th Python in Science Conference. Pasadena, CA USA; 2008. p. 11 – 15.

